# Patterns and drivers of cranial evolution in rhinos (Rhinocerotoidea; Perissodactyla): allometry, phylogeny and tempo of evolution

**DOI:** 10.64898/2026.07.25.740338

**Authors:** Mihajlo Milić, Michael Matschiner, Naomi De Leo, Gertrud E. Rössner, Davide Tamagnini

## Abstract

Despite their currently scarce taxonomic diversity, rhinocerotoids were among the most evolutionarily successful clades of large herbivorous mammals throughout the Caenozoic. Within an array of morphological adaptations, the cranial horn represents the clade’s most remarkable and name-giving feature, imposing significant morpho-functional demands on the skull. In this paper, patterns and drivers of the rhinos’ cranial evolution were investigated for the first time by compiling a three-dimensional geometric morphometric dataset of living and extinct species. Multiple aspects of morphological evolution were explored using phylogenetic comparative methods, including craniofacial evolutionary allometry (CREA – tendency of larger species to have longer faces), phylogenetic constraints, and tempo of evolution. Cranial shape variation linked to horn presence and size evolved under a phylogenetically constrained framework, with a notable transition from hornless to horned species. Rhinos significantly deviated from CREA, likely due to diversity of cranial forms and proportions that evolved independently across the clade in response to varying horn morphologies and dietary habits. Hornless and horned rhinos exhibited similar rates of cranial evolution, and shape variation was obtained through multiple episodes of accelerated evolution. This highlights the role of morphological innovations and Caenozoic global cooling events in the emergence of phenotypic diversity.

## 1. Introduction

Evolution of exaggerated or peculiar traits has been shown to significantly contribute to phenotypic diversification across the animal kingdom [1–3], highlighting their role in the dynamics and direction (i.e., tempo and mode) of morphological evolution [4–6]. Emergence of such novelties is often coupled with increased rates of evolution associated with the occupation of new, previously unexplored, ecological niches within the adaptive landscape [4–7]. In some cases, the development of extreme traits can lead to the reorganisation of complex morphological structures, such as the mammalian cranium [8,9], which can result in deviation from common evolutionary trajectories [10,11]. In comparative studies of such patterns, accounting for the fossil record is crucial for understanding past evolutionary processes that have shaped morphological diversity throughout time and space. With the dimension of geological time, fossils fill the temporal gaps and provide insight into intermediate or transitional forms [5,12,13]. This is of particular importance in clades with a small number of living species that comprise only a fraction of the total past and present species richness [14,15].

The evolution of mammalian cranial diversity is driven by various ecological, phylogenetic, and developmental factors that interact together to shape the pattern of craniofacial evolutionary allometry (CREA), which emerges as one of the most influential among mammals [16–19]. This common pattern predicts that, within clades of closely related species, larger mammals tend to have a proportionally larger or more gracile facial (i.e., rostral) part of the cranium, while the braincase appears relatively smaller [16–18]. The pattern predicted by CREA has been confirmed in numerous mammal clades, spanning a wide range of taxonomic diversity of both marsupials and placentals and encompassing a variety of different ecomorphs [16–18,20–25]. However, as for most biological rules, exceptions exist. For instance, it has been shown that sabre-toothed cats deviate from this pattern, and thus it is believed that peculiar clade-specific morpho-functional adaptations can break this evolutionary trend (e.g., biomechanical constraints related to the presence of elongated canines [10,18,20]). Different mechanisms have been proposed for explaining facial elongation in mammals [16,17]; however, Mitchell *et al.* [18] recently highlighted the importance of relaxed bite force requirements imposed by an increase in absolute size. This reasoning is best applied to carnivorous mammals, which are in general under high selective pressure for increased bite force, whereas processes and mechanisms leading to the CREA pattern in large herbivores remain less understood (but see [23,26,27]).

Originating during the early Eocene of Asia, Rhinocerotoidea (often referred to just as “rhinos”) comprises the most taxonomically and morphologically diverse group of Perissodactyla [28–30]. This clade consists of several families that diversified through the Oligocene and Miocene epochs, establishing themselves as one of the most dominant groups of large herbivores at that time [29,31]. Ranging from dog-sized *Hyracodon* (Hyracodontidae) through hippo-like forms within Amynodontidae to the largest known terrestrial mammals (Paraceratheriidae), this clade adapted to various ecological niches and habitats, accompanied by a series of morphological adaptations like derived molar dentition, reduced and specialised incisors, and diverse appendicular adaptations reflecting different modes of locomotion [29,32–36]. Nevertheless, the most evolutionarily successful rhinocerotoid clade is Rhinocerotidae (the so-called “true rhinos”), the only one to persist throughout the Neogene and Quaternary [31,37,38]. Despite the current sparse taxonomic diversity of only five extant species [39], all being highly threatened by extinction [40], rhinocerotids were an exceptionally diverse clade throughout their evolutionary history [31]. Besides their derived dental and skeletal traits (e.g., increased hypsodonty, specialised molars, shorter skulls and limbs), the appearance of keratinous cranial horns in some taxa represents the most remarkable trait of true rhinos, a feature that evolved several times independently across the phylogeny [36–38]. Horns came in different forms, numbers, and sizes, including, for instance, a gigantic frontal horn present in *Elasmotherium*, paired nasal horns in *Menoceras* and *Diceratherium*, or one larger nasal and the second smaller frontal horn characteristic of the extant black and white rhinos [31]. However, many rhinocerotids lacked horns, including the majority of basal species and tribe Aceratheriini (see [41,42]). The presence/absence of the horn, along with its form, notably influence cranial morphology in rhinos, leading to clade- and species-specific adaptations [29,36,42]. However, despite their exceptional diversity, patterns of cranial evolution in this group are poorly known. Past studies of rhino cranial evolution mostly included extant species, occasionally with a very small number of extinct species that encompass a limited geological age [43,44], and relied on a limited pool of relatively outdated morphometric analyses, with a poor phylogenetic background.

In this paper, we compile a three-dimensional (3D) geometric morphometric dataset and implement phylogenetic comparative methods with the goal of exploring different aspects of cranial diversification in rhinos, with a special focus on CREA. Most of the studies exploring CREA focus only on extant species diversity, while those including fossil data of extinct species are still rare [10,24,26,45], although it has been suggested that inclusion of paleontological data can make a significant contribution to the understanding of this pattern [20,45]. Here, we include a wide range of cranial morphologies across Rhinocerotidae in the analysis for the first time, spanning approximately 35 million years of evolutionary history. We investigate how peculiar adaptations, in this case horn presence and size, influence general morphological variation in rhinos and whether they can disrupt a pattern predicted by CREA. Horned rhinos exhibit substantial variation in the rostral part of the cranium [29], mostly linked to the type and size of their horns. This variation could reflect an array of evolutionary pathways in which different groups have developed distinct cranial morphologies, fitting different allometric trajectories throughout their evolution, to accommodate their horns. This series of morphological changes could significantly alter cranial proportions, and thus, we predict that horned rhinos could appear as an exception to CREA, while we expect hornless rhinos to follow common craniofacial allometric scaling. In addition, concordance between morphological variation and phylogenetic divergence was tested using several different approaches. The tempo of evolution was evaluated by estimating branch-specific rates across the phylogeny and comparing differences in evolutionary rates between horned and hornless rhino species. Considering the aforementioned diversity of cranial forms in horned rhinos, we hypothesise that they have higher rates of evolution compared to their hornless relatives. Since there is no broadly accepted consensus phylogeny of Rhinocerotoidea that encompasses our entire dataset, we construct one of the first total-evidence phylogenies of rhinos (along with Borrani *et al.* [46]) using published mitochondrial genomes and morphological data in order to apply phylogenetic comparative methods in our analytical framework.

## 2. Material and methods

### 2.1. Phylogenetic analysis

To infer phylogenetic relationships and divergence times of Rhinocerotoidea, total-evidence tip-dating analysis was performed using the fossilised birth-death (FBD) model implemented in BEAST 2 v.2.7.7 [47,48]. This way, we could compile a phylogeny that contains all species included in the analysis of cranial morphology (see below). The FBD model differs from other inference models in that it explicitly accounts for data on fossil occurrences [49]. As a result, higher probabilities are assigned to phylogenies that align better with the fossil record. The FBD model includes diversification and sampling processes to describe a clade’s evolutionary history through speciation and extinction dynamics [50,51]. Parameters specific to the FBD process include: speciation rate *λ*, extinction rate *μ*, fossil sampling rate *ψ* (i.e., a rate at which fossils are sampled across the tree), origin time *ϕ* (i.e., an origin of the ancestral stem lineage), and extant sampling probability *ρ* (i.e., the probability for an extant species to be included) [50,52]. Free parameters (*λ, μ, ψ, ϕ*) were estimated during the analysis, while the sampling probability of extant taxa (on which the FBD process was conditioned) was fixed to *ρ* = 1.0, as all living rhino species were included. Estimates of diversification rate (*d* = *λ*−*μ*) were drawn from an exponential prior distribution with a mean value of 1.0. The turnover parameter (*r* = *μ*/*λ*) was modelled using a uniform prior distribution with values ranging from 0.0 to 1.0. A beta prior distribution, with *α* and *β* equal to 2.0, was placed to estimate the sampling proportion (*s* = *ψ*/(*μ*+*ψ*)). An upper boundary of the origin time was set to 100 million years ago (Mya), and the lower limit to 55 Mya, modelled using a uniform prior distribution.

For the morphological data partition, a matrix of 295 discrete characters for 63 species was compiled from published studies [41,53–61] (Table S1). The scoring scheme for characters 1-282 was retrieved from Antoine [62], while characters 283-295 represented 13 newly described ones in Antoine *et al.* [41]. Morphological character states for *Diceros praecox* were scored based on images and descriptions published in Geraads [63], Geraads [64] and Guérin [65], and based on the skull specimen (KNMER 5555) available in the African Fossils database (https://africanfossils.org/). The character matrix was edited where needed, for instance, to adjust the order of characters or to align them with the scoring schemes from Antoine [62], if they were coded differently in two matrices (e.g., one character was coded as 0 in one matrix but 1 in the other). Since the number of observed states per character varied between two and four, the matrix was separated into 3 partitions (two, three, and four-state characters) to account for different rates of site evolution estimated using the Lewis Mk model [66]. This model represents a generalisation of the Jukes-Cantor matrix [67], which assumes equal probability of the change from one character state to any other. For each partition separately, a gamma distribution with four categories was used to account for rate heterogeneity among sites. The Optimised Relaxed Clock (ORC) was assumed to model rate variation of character change across lineages (i.e., branches), using a prior gamma distribution with a mean value of 10.0 [68].

Molecular sequence data of nine published mitochondrial genomes [69–72] were retrieved from GenBank (Table S1), representing all five extant and three extinct rhino species, as well as *Tapirus terrestris* serving as the outgroup. Sequences were initially aligned using MAFFT v.7.526 [73] and then separated into 13 protein-coding and two rRNA genes using AliView v.1.30 [74], resulting in an alignment of 13976 base pairs (bp). To account for potentially different rates of evolution, nucleotide sequences were concatenated to form four partitions: 1) the first codon position, 2) the second codon position, and 3) the third codon position of protein-coding genes, and 4) the 2 rRNA genes. The bModelTest package was used for each partition separately to estimate the most appropriate substitution model, rate heterogeneity across sites, and the proportion of invariable positions [75]. ORC with a prior exponential distribution (mean = 10.0) was used to model the variation in substitution rates across branches.

Fossil ages were primarily obtained from the New and Old Worlds (NOW) Database of fossil mammals [76], supplemented with the data from the relevant literature [41,69,77] and the Paleobiology Database [78]. Tip ages were specified based on the most recent occurrence of each fossil specimen. Additionally, fossil age uncertainty was considered by specifying the maximum and minimum estimated ages (of the same most recent specimens) as the upper and lower boundaries of a uniform prior distribution on the tip age. Based on estimates from Bai *et al.* [28] and Liu *et al.* [39], the root age was constrained to approximately 65-55 Mya using a log-normal prior distribution with a mean value of 60 Mya and a 0.7 Myr standard deviation (5% quantile = 56.2; 95% quantile = 67.4). A topological constraint was applied so that Rhinocerotidae and Paraceratheriidae appear as sister clades, in line with the results from Bai *et al.* [28] and Deng *et al.* [79]. Based on preliminary analysis, *Rhinoceros* spp., *Dicerorhinus* spp., *Pliorhinus* spp., *Nesorhinus* spp., *Stephanorhinus* spp., *Coelodonta antiquitatis*, and *Dihoplus schleiermacheri* were constrained to form a monophyletic clade, so that the relationships among extant taxa aligned with a recently published phylogeny based on whole-genome data [39].

The analysis was performed in three independent replicates using the Markov Chain Monte Carlo (MCMC) algorithm in the Bayesian framework implemented in BEAST 2. Trees were sampled every 6000 MCMC steps over a total of 150 million generations. Chain convergence was evaluated by comparing parameter traces and posterior marginal densities among replicates, and by assessing effective sample size (ESS) in Tracer v.1.7.2 [80]. Only one parameter from each run had an ESS below 200 (but above 100), while all parameters had ESS > 200 after combining traces across analysis replicates. Each replicate resulted in a total of 25000 trees that were combined using LogCombiner. A maximum clade credibility (MCC) tree with node heights set to median ages was produced from the combined file using TreeAnnotator, with 10% of each chain considered as burn-in. The MCC tree was subsequently pruned to match the taxon set with cranial morphology data.

### 2.2. Cranial sample and metadata collection

#### 2.2.1. Species categorisation and horn data collection

Since the taxonomy of rhinos represents a dynamic field of research [39,41,42,46,53,62], we here categorise species within “groups” according to type genus, rather than specific hierarchical taxonomic ranks (subfamily, for instance), by subdividing them into five different categories: 1) basal Rhinocerotoidea, 2) basal Rhinocerotidae, 3) elasmotheres, 4) aceratheres and teleoceres, and 5) rhinoceroses. Basal Rhinocerotoidea include Stem Rhinocerotoidea, non-rhinocerotid species from Hyracodontidae, Amynodontidae, and Paraceratheriidae. Basal Rhinocerotidae encompass basal Stem Rhinocerotidae, species without higher hierarchical clade allocation, which diverged during early stages of the true rhinos’ evolution. Elasmotheres include fossil rhinos known from the Oligocene to the Late Pleistocene, showing variable body size and generally possessing horns with great variation in their size and position [31,62]. Aceratheres and teleoceres comprise a group of extinct, mostly hornless rhinos, with some species having small nasal horns [31,42]. These two groups dominated throughout the Miocene, and recent phylogenetic studies question their monophyly and indicate a closer relationship than previously thought [42,46]. Rhinoceroses originated during the Miocene and became the most dominant group during the Pliocene and Pleistocene, eventually giving rise to all of today’s extant species [31]. All members of this group have a nasal horn, with some also developing a smaller frontal horn.

Data on horn presence and size, used in downstream analyses, were taken from published studies [31,38,42,60,81–85]. Based on these data, species were categorised into three separate groups: 1) hornless, 2) small-horned and 3) large-horned rhinos. Since keratinous horns do not preserve in the fossil record, this classification was based on authors’ descriptions of cranial specimens for extinct species, where larger horns are assumed to be associated with the more robust nasal bones with a broad base for the horn attachment. In extant species, horn size was determined based on horn measurements from published studies and the most common terminology found in the literature (large or small horn). Species possessing two horns (e.g., black and white rhino) were categorised according to the size of the larger, nasal horn.

#### 2.2.2. Cranial sample and geometric morphometrics

The dataset used for geometric morphometric analysis consisted of 33 crania, encompassing all five living rhino species and 23 fossil taxa belonging to four different families (Hyracodontidae, Amynodontidae, Paraceratheriidae, Rhinocerotidae). The cranial data sample contained all major Rhinocerotidae groups, including elasmotheres, aceratheres, teleoceres and rhinoceroses, along with several stem species. All specimens used in analyses were adults, as determined by the fully developed dentation and fusion of cranial sutures. A detailed list of specimens used in this study can be found in Table S2. Using a single or a few specimens per species has been shown to be a valid approach in macroevolutionary studies involving substantial interspecific differences [86,87]. When information from multiple specimens per species were available, morphological data were averaged to obtain species-level means, which were used in further statistical analyses. The collection of 3D cranial models consisted of scans produced by the authors using either an Artec Spider surface scanner or photogrammetry, following a digitisation protocol specifically designed for euungulate crania [88]. This sample was also integrated with 3D models obtained from online data repositories (Phenome10K, 3Dthèque MNHN - Paris, Sketchfab, and African Fossils) or from the literature (e.g., Geraads *et al*. [61]). The utilisation of datasets comprising 3D models derived from different reconstruction techniques has been shown to have a negligible effect on morphological variation and parameter estimation in mammalian macroevolutionary analysis [23,89].

Taphonomical deformations of fossil specimens, arising due to geological processes such as sediment compression, were removed from seven skull scans using the protocol described by Schlager *et al.* [90], which is implemented in the *Morpho* R package [91]. Landmark configuration used to describe cranial morphology included 25 fixed anatomical landmarks and 17 semi-landmarks forming a curve along the roof of the skull (Figure S1). Definitions of landmark positions are given in Table S3. To avoid inter-operator biases, landmark digitisation was performed by the same person (M. Milić) using the Stratovan Checkpoint software (v. 2025.05.23) [92]. To evaluate the effect of digitising error (i.e., precision and repeatability of landmark configuration), landmarks were digitised twice for each specimen within two separate time intervals, using a subsample of 25 fixed landmarks previously defined (i.e., the same landmark configuration just without semilandmarks). Based on the linear model implemented in the *bilat.symmetry()* function from the *geomorph* R package [93,94], digitising error accounted for a minor fraction of the total shape variation (R^2^ = 0.85%). Additionally, a cluster dendrogram was built for cranial shape data using the unweighted pair group method with arithmetic mean (UPGMA) applied to the Procrustes distance matrix among specimens. The first and second replicates of each specimen clustered together in all of the cases, indicating high precision and repeatability of the landmark configuration. To evaluate potential error in estimating skull size, a correlation between centroid size values for the two sets of replicates was calculated. A high correlation coefficient was retrieved (r = 0.99; P < 0.001), indicating high repeatability. In cases of damaged specimens with missing skull parts, missing landmarks were primarily estimated using the *fixLMmirror()* function from the *Morpho* package. This function calculates landmark positions by the mirror reflection of their non-missing bilateral counterparts along the midplane of the skull. Unilateral missing landmarks and bilateral landmarks missing on both sides of the skull were estimated using the *estimate.missing()* function from the *geomorph* package using the thin-plate spline (TPS) method. In total, 87 missing landmarks were estimated (6.27% of total landmarks), with 50 estimates using the mirror reflection approach and 37 using the TPS method.

To remove non-shape variation arising due to differences in scale, translation and rotation, a Generalised Procrustes Analysis (GPA) was performed using the *gpagen()* function on the raw landmark coordinates, with the bending energy method for sliding semi-landmarks [95,96]. Using the *bilat.symmetry()* function, the symmetric component of shape variation was extracted and used as the shape variable in all downstream analyses [97]. In all further analyses involving size variation, the logarithm of centroid size (logCS) was used as the input variable. All statistical analyses of morphometric data were performed using the R (v. 4.5.3) programming language [98], primarily within the *geomorph* (v. 4.1.0) package, unless stated differently.

#### 2.2.3. Comparative phylogenetic methods – evolutionary patterns of cranial morphology

The presence and the strength of phylogenetic signal in size and shape data were evaluated using Blomberg’s K and K_mult_ metrics as proxies, respectively [99,100]. Phylogenetic signals were estimated using the *physignal()* function. A principal component analysis (PCA) was performed to explore cranial shape variation and relations between different horn groups within phylomorphospace, using the *gm.prcomp()* function. Following Collyer and Adams [101], phylogenetically-aligned component analysis (PaCA) was performed to assess shape variation relative to the phylogeny. This analysis aligns the shape data according to an axis of greatest phylogenetic signal rather than axis of greatest dispersion, so that the first few axes bear the strongest signal [101].

The effect of CREA (and horn variation) on cranial shape was tested using Phylogenetic Generalised Least Squares (PGLS) analysis for high-dimensional data implemented within *procD.pgls()* function [102], utilising the ***shape ∼ logCS * horn*** formula, where the “horn” was a categorical variable containing information on horn presence and size (absent, small or large). This analysis is performed under the Brownian motion model of evolution, which generally describes a process of neutral evolution where phenotypic variation increases at a constant rate through time. The significance level was estimated using a non-parametric approach with 10 000 iterations (9 999 permutations and 1 observed). Considering its exceptionally large size, *Paraceratherium grangeri* was excluded from the allometric analysis, as this species appeared as an outlier in the preliminary assessment. In addition to PGLS, Ordinary Least Squares (OLS) analysis, which does not account for phylogenetic correlation (i.e., species relatedness), was performed within the *procD.lm()* function under the same settings. Furthermore, a series of subset allometric analyses was conducted separately on the groups of hornless, small-horned and large-horned species, including both PGLS and OLS approaches.

To evaluate the mismatch between cranial shape and the phylogeny, a tanglegram was constructed using the *tanglegram()* function of the *dendextend* (v. 1.19.1) package [103]. Initially, a matrix of pairwise Procrustes distances between species was calculated using Procrustes shape coordinates, which was subsequently used to construct a cluster dendrogram with the UPGMA approach. The obtained phenogram of cranial shape was then used in a comparison with the phylogenetic tree. Differences in net rates of cranial shape evolution between horned and hornless Rhinocerotidae species were tested using the *compare.evol.rates()* function, under a Brownian motion model of evolution [104]. The significance level was estimated through 10000 iterations using the “simulation” method, in which tip data are obtained under Brownian motion using a common evolutionary rate pattern for all species on the phylogeny [105]. Additionally, branch-specific rates of cranial evolution were estimated by fitting the matrix of pairwise Procrustes distances between species into the *RRphylo()* function from the *RRphylo* (v. 3.0.2) package, which performs the phylogenetic ridge regression under a series of trait evolution models to calculate evolutionary rates [106].

## 3. Results

### 3.1. Phylogeny – fossilised birth-death analysis

Although the main goal of this paper is to explore the evolution of cranial morphologies in rhinos, our phylogenetic analysis provides valuable new insights into the evolutionary history of Rhinocerotidae (Figure 1). Therefore, the most important results from the FBD analysis are presented and discussed in this section (with systematic classification following Borrani *et al.* [46]).

**Figure 1.**
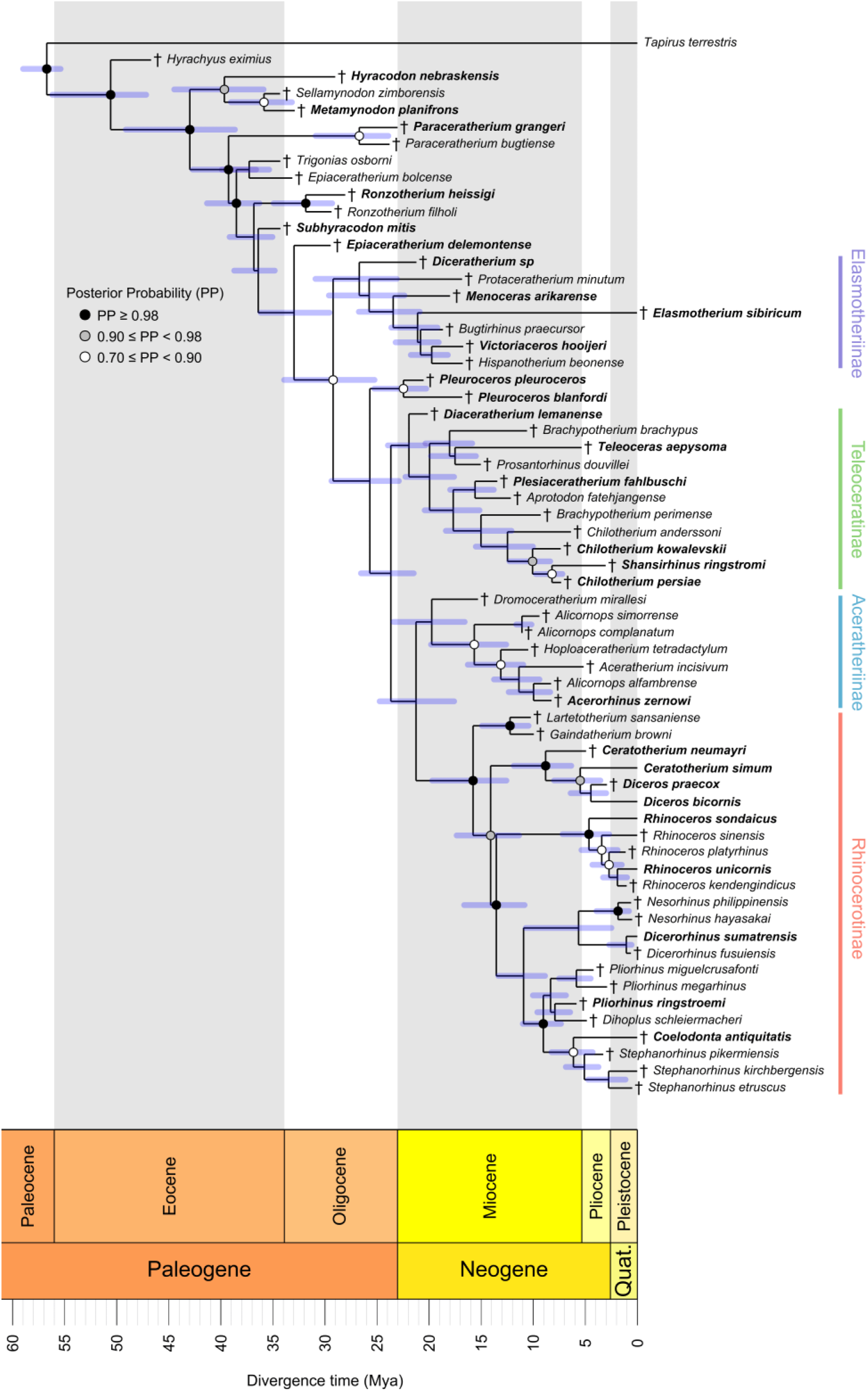
Time-calibrated phylogenetic tree of rhinos inferred with the total-evidence approach and used in the analysis of cranial morphology. Cross symbols before taxon names mark extinct species. Species in bold were included in the geometric morphometric dataset. Blue bars at nodes represent 95% highest posterior density (HPD) intervals for node ages. Tree visualisation was performed in the *strap* (v.1.6) R package [150].

Among basal Rhinocerotidae, *Epiaceratherium* was not recovered as monophyletic, since *E. bolcense* appeared as the most basal rhinocerotid together with *Trigonias osborni*, while *E. delemontense* was placed closer to the crown group. In line with the results of Borrani *et al.* [46], *Epiaceratherium* is most likely polyphyletic, consisting of multiple independent lineages, in contrast to what was previously thought [107]. The position of Elasmotheriinae has been widely debated, with a few recent studies placing it as the sister group to Rhinocerotinae [42,46], while this clade has traditionally been placed more basally [53,58,108]. The results from our analysis supported the latter hypothesis, with *Diceratherium* representing the basalmost genus within the group, although posterior probabilities within the clade were low. *Protaceratherium*, which has been classified differently in different studies [42,46,53,109], was recovered as an early-diverging member of Elasmotheriinae in our analysis, consistent with Fraser *et al*. [110].

The phylogenetic positions of Teleoceratinae and Aceratheriinae (here classified as subfamilies, but see Antoine *et al.* [41] and Antoine *et al.* [53] for alternative systematics) have been contentious, and their monophyly has recently been challenged [42,46]. Some studies have proposed Teleoceratinae as a sister clade to Rhinocerotinae, with Aceratheriinae diverging as a sister clade to the Teleoceratinae-Rhinoceratinae [41,53,59,60,111,112]. Contrarily, a few new phylogenetic studies have identified Aceratheriinae and Teleoceratinae as more closely related, with significant intermixing between clades [42,46,107]. The FBD phylogeny presented here revealed Aceratheriinae and Rhinocerotinae as sister groups, with Teleoceratinae diverging basal to them; however, these relationships were characterised by low posterior probabilities (PP < 0.50).

Congruent with Borrani *et al.* [46], *Chilotherium* and *Shansirhinus* clustered within Teleoceratinae, rather than Aceratheriinae as classified traditionally [53,54,77]. However, *Shansirhinus ringstroemi* appeared as a sister species to *Chilotherium persiae* with a moderate posterior probability (PP = 0.73), questioning the monophyly of the *Chilotherium*. *Plesiaceratherium fahlbuschi* also showed affinities towards Teleoceratinae, although it has been considered an acerathere originally [113,114]. *Brachypotherium* was not identified as monophyletic, which aligns with Borrani *et al.* [46]. In agreement with Lu *et al.* [42], *Brachypotherium brachypus* was more closely related to *Teleoceras* and *Prosantorhinus*, whereas *B*. *perimense* was a sister species to the *Chilotherium*-*Shansirhinus* clade based on our analysis.

*Dromoceratherium mirallesi* was recovered as the basalmost Aceratheriinae, as demonstrated by Borrani *et al.* [46], and opposed to previous studies where it appeared as an isolated lineage [41,53,60]. *Alicornops* appeared polyphyletic as previously noted [53], with *A*. *alfambrense* being a sister species to *Acerorhinus zernowi*, albeit with weak posterior probability (PP = 0.58). On the other hand, *Alicornops complanatum* was sampled as a direct ancestor of *Alicornops simorrense*. It should be noted that most of the nodes within Teleoceratinae and Aceratheriinae had low posterior probabilities (PP < 0.70), suggesting weak support to infer relationships among species with high certainty.

Within Rhinocerotinae, *Gaindatherium browni* and *Lartetotherium sansaniense* were identified as the most basal species of the clade with strong support (PP = 1.00), in agreement with several previous studies [42,58,60,108,110]. *Ceratotherium* was not monophyletic, since *C. neumayri* appeared as the ancestral lineage to *Diceros* spp. and *C. simum*, as previously noted by some authors [41,59,84,110]. *Diceros bicornis* and *Diceros praecox* were identified as sister species, suggesting the monophyly of the genus. Together with Borrani *et al.* [46], this is the first study that includes *D. praecox* in a phylogenetic analysis and significantly contributes to the understanding of its phylogenetic position. The position of *Nesorhinus* has been shown to be variable across studies. Some identified it as a sister genus to *Dicerorhinus* [60] and others as more closely related to *Rhinoceros* [46], whereas further studies suggested it as a basal genus in the *Rhinoceros*-*Dicerorhinus* clade [41,110,111]. The first hypothesis was supported by the present analysis, as *Nesorhinus* and *Dicerorhinus* formed a monophyletic clade (PP = 0.41). *Dihoplus schleiermacheri* appeared within *Pliorhinus* spp., highlighting the debatable taxonomy of both genera [46]. *Stephanorhinus* appeared as monophyletic, with *Coelodonta* appearing as a sister genus (PP = 0.78), which aligns with several previous molecular- and morphology-based phylogenies [39,42,46,115,116].

In summary, the presented phylogeny emphasises the complex evolutionary patterns within Rhinocerotidae that have long been debated. The relationships among major crown subfamilies have proven hard to resolve, as in earlier studies [42,46]. Moreover, we provide one of the first total-evidence phylogenies of rhinos utilising ancient mitochondrial genomes of all currently available extinct species and all extant species, which will hopefully facilitate the resolution of further questions in the field.

### 3.2. Evolutionary patterns in cranial morphology

Phylogenetic signal was significant for both cranial shape (P = 0.0001) and size (P = 0.027) data, with relatively low values of the K parameter (Kmult = 0.66 and Blomberg’s K = 0.56, respectively), suggesting that closely related species are less similar than expected under Brownian motion. In terms of variation in cranial size within Rhinocerotidae, larger species showed a tendency to evolve relatively larger horns, whereas small-horned species appeared in the range of sizes similar to the hornless rhinos (Figure S2). *Victoriaceros hooijeri* seemed to be the only exception, given its large cranial size despite having a small horn. The first two principal components (PC1 and PC2) described 51% of the total shape variation, showing a noticeable patterning with respect to the horn presence and size (Figure 2; see Figure S3a for the phylomorphospace with all taxon labels). At the most negative end of PC1, members of non-Rhinocerotidae taxa (*Paraceratherium*, *Hyracodon*, and *Metamynodon*) separated from Rhinocerotidae, emphasising the derived cranial shape of “true rhinos”. The most negative PC1 values were found for specimens characterised by a more convex or flatter skull roof, along with less pronounced and robust nasal bones (Figure 2 and Figure S4a). This included several Oligocene representatives such as *Paraceratherium grangeri* (Paraceratheriidae), *Metamynodon planifrons* (Amynodontidae), and *Epiaceratherium delemontense* (a stem rhinocerotid). At the positive end of PC1 appeared the species with a more concave skull roof reflected through a less expanded nuchal crest and generally higher posterior part of the skull, together with curved nasal bones associated with the presence of small-horned specimens such as *Victoriaceros hooijeri* and extant Asian rhinos (*Dicerorhinus sumatrensis* and *Rhinoceros* spp.). Small-horned species differentiated from their hornless relatives along PC1, with some specimens (e.g., *Diceratherium* sp. and *Shansirhinus ringstromi*) showing greater affinities towards the morphospace regions dominated by hornless representatives. The small-horned group seemed to bridge between hornless and large-horned species to a certain extent, with Miocene taxa such as *Menoceras arikarense*, *Diceratherium* sp. and *Pleuroceros pleuroceros* exhibiting transitional cranial shapes. Large-horned species occupied a distinct area of the phylomorphospace associated with the positive end of PC2, which was dominated by Neogene and Quaternary specimens such as *Coelodonta antiquitatis*, *Diceros praecox,* and *Pliorhinus ringstroemi,* characterised by dolichocephalic crania with robust nasal bones, slender zygomatic arches and relatively less expanded cheek tooth row (Figure 2 and Figure S4b). *Elasmotherium sibiricum* showed the greatest divergence relative to its closely related species within the phylomorphospace, which was driven by the presence of a huge hemispheric frontal prominence, a unique cranial characteristic in our dataset. The negative end of PC2 was primarily occupied by hornless specimens that have a more brachycephalic cranial shape with high, robust zygomatic arches, less pronounced nasals, and an expanded cheek tooth row associated with a large molar dentition. This region included Miocene and Pliocene taxa such as *Teleoceras aepysoma*, *Chilotherium persiae*, *Shansirhinus ringstromi* and *Pleuroceros blanfordi*, together with a basal rhinocerotid *Ronzotherium heissigi*. Phylogenetically-aligned component analysis (PaCA) showed a generally similar distribution of taxa and groups as the phylomorphospace analysis, implying that diversification of cranial shape in relation to the horn variation may be coupled with phylogenetic divergence (Figure S3b). The negative end of the first component was occupied by non-Rhinocerotidae taxa, along with a few basal rhinocerotids, while at the more positive values, the transition from hornless through small-horned to large-horned rhinos was generally evident along the second component. Evaluation of phylogenetic signal (Blomberg’s K) through components revealed that the greatest amount of signal was concentrated in the first few components for both PCA and PaCA (Figure S3b), supporting the premise that cranial shape variation across different horn groups is correlated with phylogenetic diversification.

**Figure 2.**
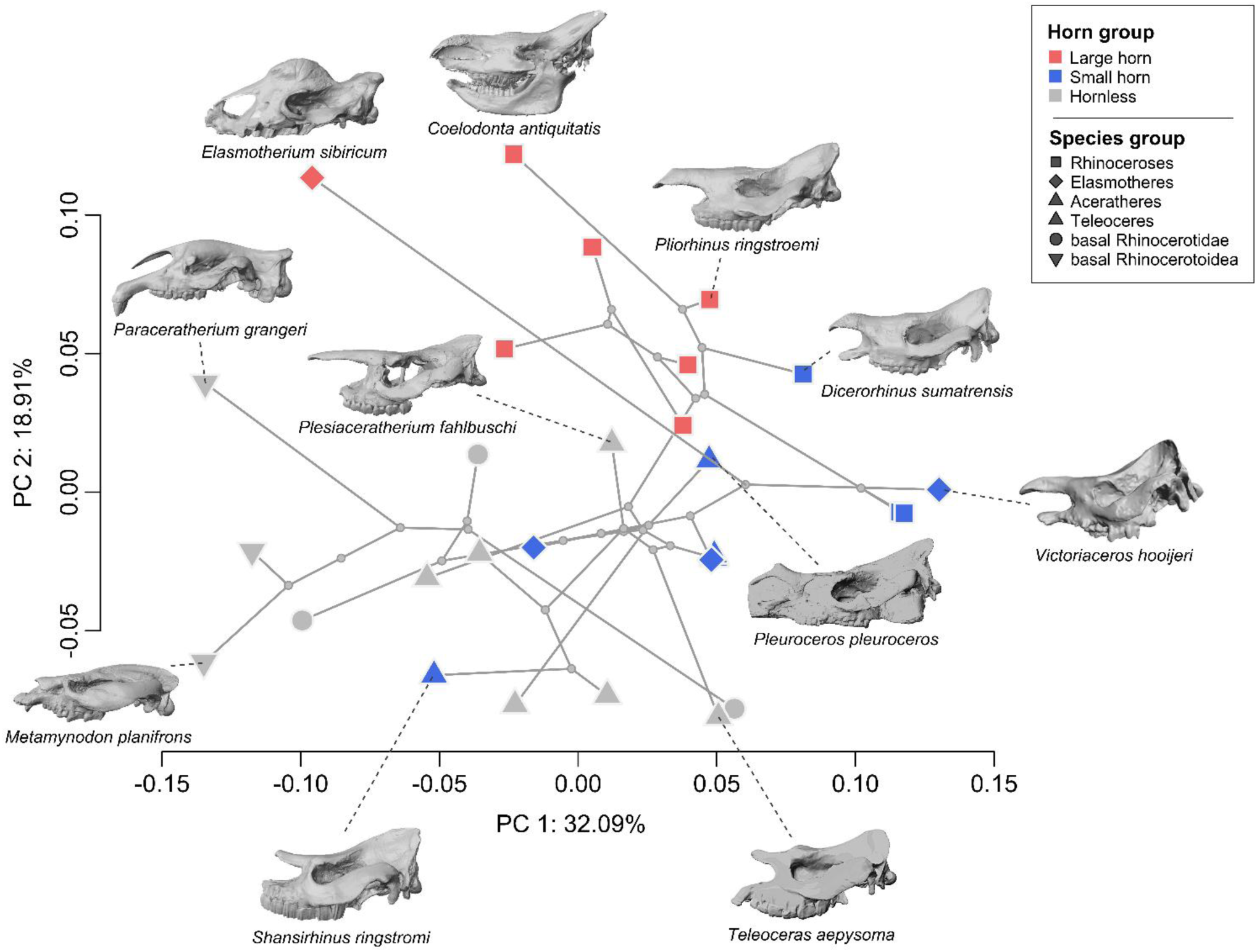
Phylomorphospace defined by the first two principal components of cranial shape across Rhinocerotoidea. The symbols represent species groups, and the colours indicate groups based on horn presence and size. Skulls from the lateral view are shown to illustrate the diversity of cranial forms. See Figure S3a for the representation with all taxon labels and Figure S4 for shape changes at the positive and negative extremes of the two PCs.

PGLS analysis for the full dataset revealed statistically non-significant effects of evolutionary allometry, horn variation and their interaction on cranial shape (P > 0.05 in all cases; Table 1). In addition, subset PGLS regressions indicated non-significant results for all tested groups (hornless, small-horned, and large-horned; P > 0.05 in all tests). In contrast, evolutionary allometry was statistically significant in the OLS analysis of the full dataset, explaining 11.68% of the total shape variation, along with a significant *logCS × horn* interaction, suggesting divergent allometric trajectories among the three horn groups (Table 1 and Figure 3a). Allometric shape changes for the full dataset were associated with a relative reduction of the rostrum in larger species, except for the nasal region, which increased in relative height and length (Figure 3b). The changes in the posterior part of the cranium associated with increasing size were mostly related to a lower position of the zygomatic arches and expansion of the nuchal crest, overall resulting in a pattern that significantly deviates from CREA. For subset OLS regressions, allometry was significant only in large-horned rhinos, where it accounted for a large proportion of shape variation (R^2^ = 31.15%; P = 0.0308), whereas in hornless and small-horned species it appeared statistically non-significant (P > 0.05). Allometric shape changes in the large-horned group were partially compatible with the CREA (Figure 3b). Cranial gracilisation was apparent in larger species, resulting in elongation of the rostrum along the sagittal plane. However, relative expansion of the braincase was also evident with an increase in overall cranial size, contrasting the pattern predicted by CREA.

**Figure 3.**
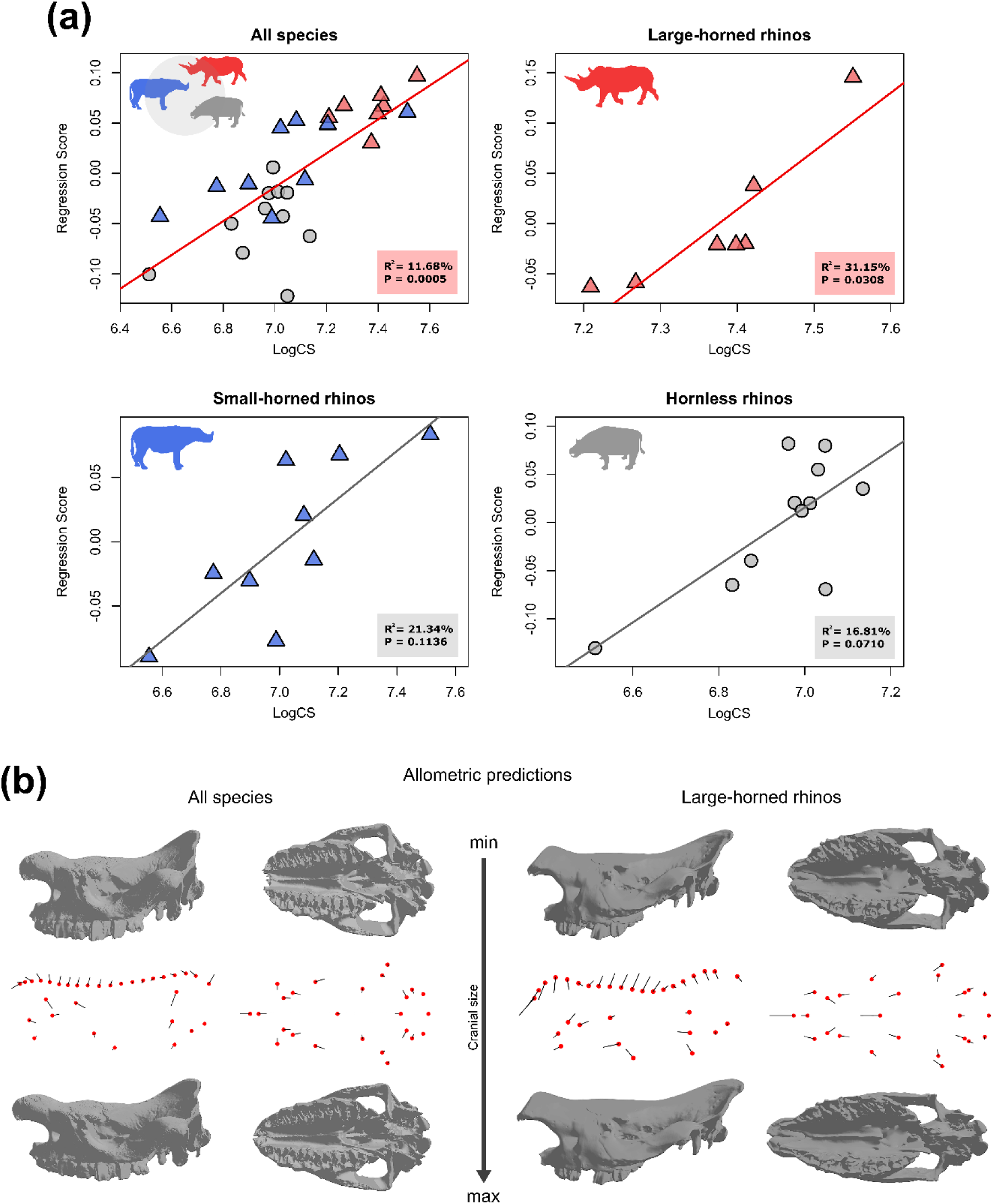
Evolutionary allometry in Rhinocerotoidea. Allometry scatterplots for the full dataset and different horn groups derived from OLS regressions of cranial shape against the logarithm of centroid size (a). Allometric shape changes represented as transitions from the smallest to the largest specimen for the full dataset and large-horned rhinos (b). Skull images represent warped 3D surfaces. Rhino silhouette icons were obtained from PhyloPic.

**Table 1.** Effect of size (logCS) and horn variation on cranial shape in rhinos. Separate regressions were performed for each sample group.

| Method | Sample | Factor | Df | R<sup>2</sup> (%) | F | Z | P |
| --- | --- | --- | --- | --- | --- | --- | --- |
| PGLS | All species | logCS | 1 | 4.77 | 1.28 | 0.46 | 0.3374 |
|  |  | horn | 2 | 11.18 | 1.50 | 0.79 | 0.2285 |
|  |  | logCS × horn | 2 | 5.50 | 0.74 | -0.04 | 0.5131 |
|  | Large-horned | logCS | 1 | 16.09 | 0.96 | 0.33 | 0.3894 |
|  | Small-horned | logCS | 1 | 13.66 | 1.11 | 0.46 | 0.3541 |
|  | Hornless | logCS | 1 | 6.94 | 0.67 | 0.07 | 0.4672 |
| OLS | All species | logCS | 1 | 11.68 | 4.23 | 3.11 | **0.0005** |
|  |  | horn | 2 | 20.11 | 3.64 | 3.52 | **0.0001** |
|  |  | logCS × horn | 2 | 10.15 | 1.84 | 1.79 | **0.0353** |
|  | Large-horned | logCS | 1 | 31.15 | 2.26 | 1.59 | **0.0308** |
|  | Small-horned | logCS | 1 | 21.34 | 1.90 | 1.29 | 0.1136 |
|  | Hornless | logCS | 1 | 16.81 | 1.82 | 1.50 | 0.0710 |

Abbreviations: logCS – logarithm of centroid size; PGLS – Phylogenetic Generalized Least Squares ; OLS – Ordinary Least Squares; Df – degrees of freedom; R<sup>2</sup> – coefficient of determination; F – F value; Z – effect size; P – significance level.

Horn variation (in both presence and size) had a significant effect on cranial shape when the phylogeny was not included in the model (OLS; P = 0.0001; Table 1), explaining 20.11% of shape variation. Generally, a significant decrease in R^2^ (coefficient of determination) and Z (effect size), and an increase in P values can be noted for the PGLS models in comparison with the OLS, suggesting that cranial shape diversification is probably constrained by phylogeny, in line with observations above.

Despite significant phylogenetic signal for cranial shape, clustering analysis indicated a generally high mismatch between phylogenetic relationship and the shape dendrogram, especially in basal Rhinocerotidae and elasmotheres (Figure 4), potentially suggesting a certain degree of morphological convergence. *Elasmotherium sibiricum* split first from the shape phenogram, emphasising its distinctive cranial morphology compared to other species included in the dataset. Basal Rhinocerotoidea clustered together, mostly in correspondence with phylogeny, whereas hornless and small-horned rhinocerotids appeared largely intermixed. For instance, a few hornless and small-horned aceratheres showed a basal position in the phenogram, clustering together with an early rhinocerotid *Epiaceratherium delemontense*, possibly highlighting their underived cranial shape. Compared to other groups, extant rhinos and their relatives showed higher phylogenetic inertia, splitting into two clusters, with one encompassing the majority of large-horned species, including *Diceros* spp. and *Ceratotherium* spp. together with the woolly rhino (*Coelodonta antiquitatis*) as an outer branch. The second group comprised extant Asian rhinos and *Pliorhinus ringstroemi*, mixed with small-horned elasmothere *Victoriaceros hooijeri* and hornless acerathere *Plesiaceratherium fahlbuschi*. Most of the species in this cluster share relatively long nasal bones, which likely contributed to their grouping, although it includes species with different horn forms and variable morphology of the nasals. *Victoriaceros hooijeri* showed a close relation to two *Rhinoceros* species, as previously indicated by the phylomorphospace analysis (see Figure S3a), whereas *Plesiaceratherium fahlbuschi* grouped with *Dicerorhinus sumatrensis* and *Pliorhinus ringstroemi.* One of the groups in the shape phenogram included most of the brachycephalic species with broad and thick zygomatic arches (e.g., *Menoceras arikarense*, *Chilotherium persiae*, *Teleoceras aepysoma*, etc.), suggesting that brachycephaly represents a derived trait in rhinos, with a significant role in the structuring of cranial shape variation.

**Figure 4.**
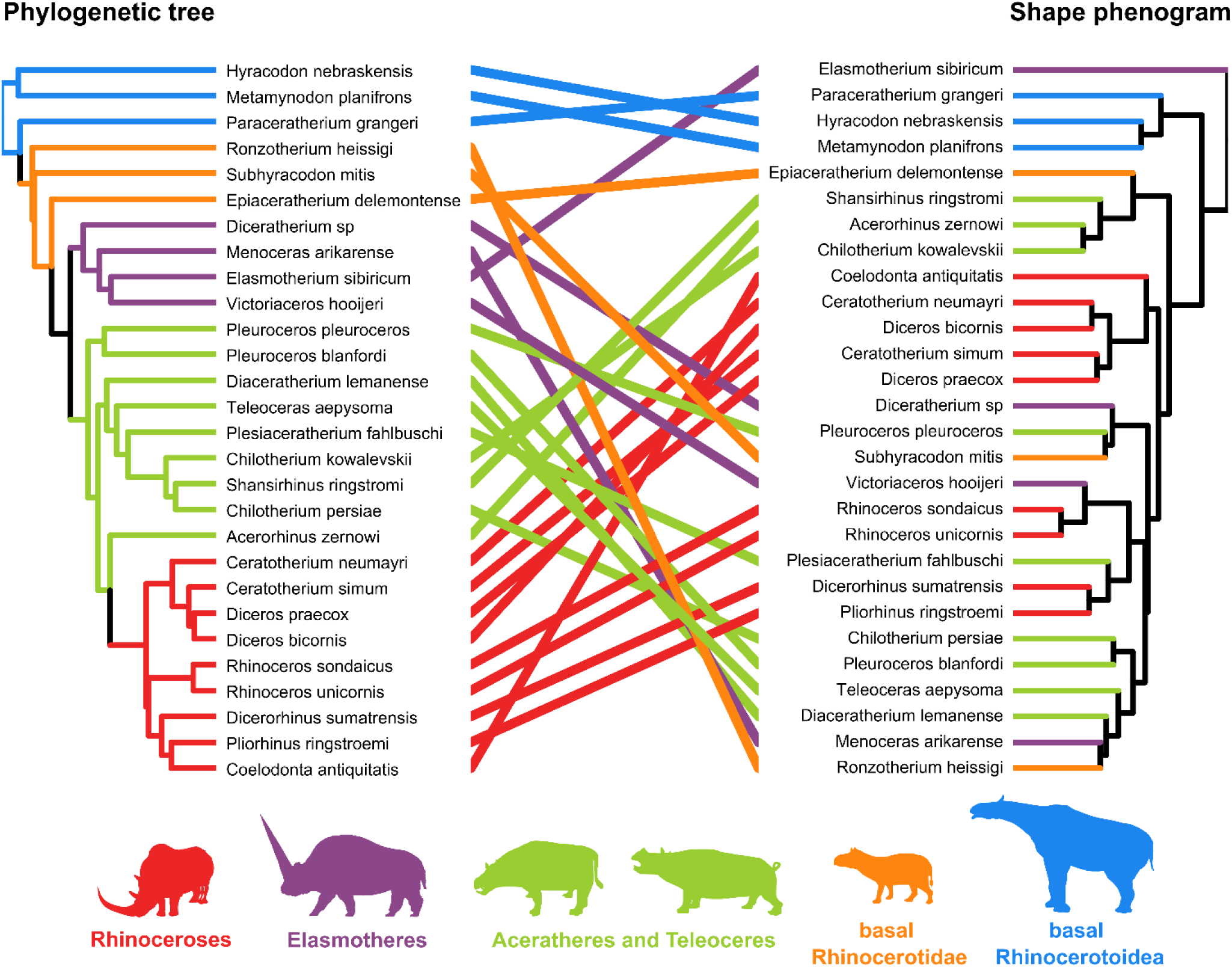
Tanglegram showing a mismatch between the phylogenetic tree and the skull shape phenogram derived from the Procrustes distance matrix among specimens. Rhino silhouette icons were obtained from PhyloPic.

Differences in net rates of cranial shape evolution were statistically non-significant (P = 0.3622; Z = 0.41) between hornless (net rate: 1.42 × 10^-5^) and horned (net rate: 1.16 × 10^-5^) rhinocerotids, with a ratio of 1.23. However, changes in cranial shape throughout the evolutionary history of Rhinocerotidae did not occur gradually, as indicated by a series of events of increased evolutionary rates across the phylogeny (Figure 5). At the early stage of Rhinocerotidae evolution, a high rate of morphological change occurred at the branch that splits from *Subhyracodon mitis*, leading to the crown groups. Moderately high rates were estimated for the branches representing origins of major rhinocerotid clades, including elasmotheres and teleoceres together with certain aceratheres (clade encompassing *Diaceratherium lemanense* to *Shansirhinus ringstromi*). A rapid burst occurred at the ancestral lineage of hornless *Shansirhinus ringstromi* and *Chilotherium persiae*, and multiple acceleration phases took place during the evolutionary history of the rhinoceroses (extant rhinos and their relatives). The first phase occurs at the origin of the clade leading to extant African rhinos and their relatives (genera *Diceros* and *Ceratotherium*), subsequently followed by a high increase in rate at the branch encompassing *Diceros* spp. and *C. simum*. Additionally, a high rate of morphological evolution was inferred for the ancestral lineage of *Pliorhinus ringstroemi* and *Coelodonta antiquitatis*, along with moderately high rates for *Rhinoceros sondaicus* and *Coelodonta antiquitatis*.

**Figure 5.**
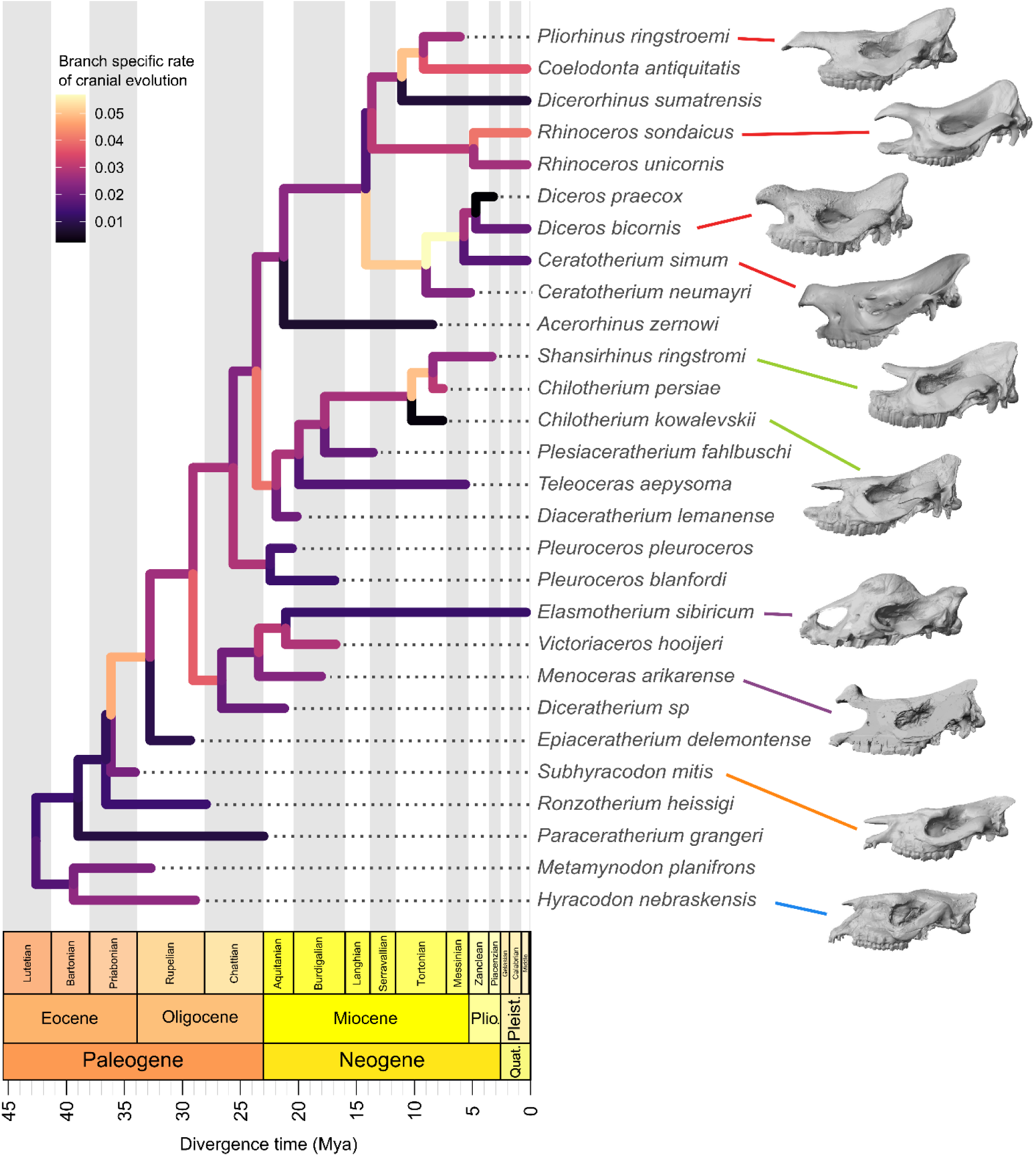
Branch-specific rates of cranial evolution across the phylogeny of Rhinocerotoidea. Note a series of acceleration phases during early-stage rhinocerotid evolution and at the origin of major groups within Rhinocerotidae, as well as within Rhinocerotinae, the clade formed by extant rhinos and their extinct relatives. Abbreviations: Plio. – Pliocene; Pleist. – Pleistocene; Quat. – Quaternary.

## 4. Discussion

Rhinoceroses represent one of the most taxonomically and morphologically rich perissodactyl groups; nonetheless, patterns and drivers of this diversity remain largely understudied compared to other clades of large herbivores such as equids and artiodactyls. Here, cranial variation in Rhinocerotidae spanning their entire evolutionary history (∼ 35 million years) was studied by compiling and analysing a 3D morphometric dataset including, for the first time, specimens from all major subgroups to explore the role of peculiar adaptations (i.e., the presence of the horn) in morphological diversification. In addition, we present one of the first total-evidence phylogenies of Rhinocerotidae, using mitochondrial genomes from both extinct and extant rhinos within a fossilised birth-death framework, demonstrating the model’s utility in comparative studies like this.

### 4.1. Effect of horn diversification on cranial morphology

A diversity of cranial forms was revealed by the phylomorphospace analysis (Figure 2), showing notable patterning with respect to horn variation, as some small-horned representatives and a hornless *Plesiaceratherium fahlbuschi* bridged between hornless and large-horned rhinos. This transition was primarily reflected through the morphology of the nasal bones, which generally appear longer and more robust with the development of horns. However, certain small-horned rhinos occupied distinctive regions of the morphospace (e.g., *Rhinoceros* spp. and *Victoriaceros hooijeri*) or have grouped with their hornless relatives (*Shansirhinus ringstromi*), deviating from the transitional forms. Since these species are part of more advanced lineages within specific rhino clades, this divergence could likely be a consequence of the acquisition of derived cranial features associated with a more phylogenetically constrained morphotype. Therefore, small-horned rhinos with a more phylogenetically basal position could provide valuable insight into gradual change in cranial morphology adapted to accommodate nasal horns.

Despite the significant contribution of horn presence and size to the composition of cranial shape variation, this pattern was largely phylogenetically constrained, suggesting that it was more a consequence of shared ancestry than explicit ecomorphological divergence. This is not surprising, considering that the horn presence is mostly limited to two rhinocerotid clades, elasmotheres (Elasmotheriinae) and rhinoceroses (Rhinocerotinae according to Borrani *et al.* [46]). Almost all representatives of these two clades possess/ed horns, while the remaining lineages were mostly hornless, with a few exceptions [31]. However, ascribing the observed variation solely to shared ancestry would be misleading, as horned rhinos exhibit a suite of morpho-functional cranial adaptations that reflect their ecology [43]. This was best exemplified by the distinctive cranial shape of *Elasmotherium sibiricum*, with a large frontal prominence serving as support either for a long horn or a massive dome [31,117], resulting in the outlier appearance in analyses above (Figures 2 and 4). Interestingly, only larger species tend to evolve relatively larger horns based on our dataset, which may imply specific biomechanical pre-requirements in order to evolve such an exaggerated trait. Perhaps species with larger crania develop stronger neck musculature, associated with an expanded occipital plane, to support the weight of large horns, which would be consistent with limited evidence from previous intraspecific and intrageneric studies [118,119]. Thus, if true, to evolve large horns, species would initially have to exhibit sufficient cranial size, accompanied by adequate musculature, suggesting a constrained biomechanical framework of morphological change. This hypothesis may be supported by the example of the small-horned *Victoriaceros hooijeri* that possessed a large cranium (Figure S2), while its sister species, *Victoriaceros kenyensis*, showed similar cranial size but developed large horns [61,120], possibly indicating that this genus reached sufficient cranial size to subsequently evolve increased horn size in one of its species. Further studies should focus on biomechanical aspects of horn development to better understand this relationship and the evolution of pronounced horn size in rhinos.

### 4.2. Craniofacial evolutionary allometry

Rhinoceroses embody a valuable model for studying evolutionary allometry due to their peculiar rostral adaptations and the diversity of cranial forms, as shown by the present analysis. A spectrum of shapes that display different craniofacial proportions was revealed, ranging from brachycephalic *Teleoceras* to dolichocephalic *Coleodonta*. In agreement with the initial hypothesis, rhinos generally appeared as an exception to the common pattern of evolutionary allometry within mammals. The allometric effect was insignificant in analyses with phylogenetic comparative methods (PCMs), whereas the allometry was impactful when phylogenetic relatedness was not accounted for. This methodological pattern was already addressed by Mitchell *et al.* [18], who suggest that evolutionary allometry may be obscured when there is substantial covariation between size and phylogeny, that is, when size data bear significant phylogenetic signal (as in our case). In such cases, authors indicate that use of PCMs may not be necessary to test the correlation between size and shape [18]. However, Wisniewski and Slater [26] argued against this interpretation and emphasised the importance of incorporating phylogenetic data in the analysis of CREA. They also empirically demonstrated that correlations between traits are more reliably inferred when using PCMs compared to standard regressions, in which evolutionary patterns can be overlooked [26]. It should be mentioned that PGLS tests used to assess CREA relied on the Brownian motion model of evolution, which may not be the best fit for our data. However, it is currently the only model implemented for strictly geometric morphometric data, and further analysis on the sensitivity of these tests could be informative. In any case, even if allometry was significant by applying OLS tests on the complete dataset, the pattern of allometric shape changes deviated from CREA predictions, with divergent allometric trajectories among horn groups. This deviation can be at least partially attributed to substantial variation in the morphology of the nasal bones of horned rhinos, since the evolutionary allometry in hornless species was non-significant, in contrast to our hypothesis. Hence, we propose several other factors that likely contributed to deviation from CREA, including ecomorphological adaptations that mostly covary with dietary preferences, independently of cranial size or phylogeny.

Throughout their evolutionary history, rhinocerotoids have evolved a series of craniodental adaptations, including reduced anterior dentition, molarisation of premolars, and increased hypsodonty [29,37,38,107]. A trend of reduced anterior dentition (incisors and canines) is often coupled with a reduction of the premaxillary bone that bears these teeth, and it is highly correlated with species’ dietary habitats [29,36,60]. For instance, some derived rhinoceroses, such as *Coelodonta* or *Rhinoceros*, have a fully developed premaxilla, resembling the ancestral state of basal rhinocerotids (e.g., *Subhyracodon* or *Epicaeratherium*), whereas in other species it is greatly reduced (e.g., *Diceros* or *Chilotherium*) or highly specialised (e.g., *Teleoceras*). The reduction of the premaxilla in larger species would thus imply a pattern opposite to CREA, as it would result in a reduced rostrum for a larger cranium. However, it should be noted that the morphology of the premaxilla was not assessed through the above-defined landmark configuration due to its rare preservation in the fossil record and the already mentioned reduction in some species. Nonetheless, changes in this character’s state throughout rhinos’ evolution are very likely an important factor contributing to deviation from CREA. Molarisation of premolars and increased hypsodonty represent general trends in rhinocerotid evolution, however, they are mostly pronounced in highly specialised grazing species such as *Elasmotherium sibiricum*, *Ceratotherium simum,* or *Shansirhinus ringstromi* [29,37,77]. In contrast, browsers and mixed feeders exhibit brachyodont, mesodont, or subhypsodont molars [118,121]. Hence, a variety of dental adaptations during the evolutionary history of rhinos might have triggered alterations of cranial proportions due to the need to house large, high-crowned teeth in grazing species, eventually contributing to deviation from CREA.

Although horned rhinos bear the greatest amount of variation in the morphology of the nasal region, hornless species also show substantial variation in the nasal length and the depth of the nasal notch [29,42]. This variation is usually correlated with feeding habitats, as browsing species (e.g., *Plesiaceratherium* spp. or *Aphelops* spp.) often have deeper nasal notches and longer nasals, associated with an attachment of muscles of a prehensile upper lip used to grasp higher foliage [29,38,122]. Therefore, differences in the morphology of the nasal region in hornless rhinos, together with the above-mentioned dental adaptations, are probably the most influential factors contributing to the shift from CREA in this group. Furthermore, rhinocerotids show significant variation in the shape of the posterior cranium, which determines the neutral head position among species [83]. As an adaptation to a grazing diet, several rhino species exhibit posteriorly elongated nuchal crests (e.g., *Ceratotherium simum* and *Coelodonta antiquitatis*), allowing a more vertical head position closer to the low vegetation, resulting in a dolichocephalic cranial shape [36,83]. Browsers, in contrast, show more horizontal head posture associated with a straight or even forward-inclined occiput and relatively more anteriorly positioned orbits. This feature likely contributed to the absence of reduced braincase in larger species, especially in large-horned rhinos, where an increase in cranial size was accompanied by an expanded posterior cranium. However, large-horned rhinos were the only ones to show any signs of CREA, since the facial elongation was apparent with increased size. Evolution of large horns represents a relatively rare event across Rhinocerotidae phylogeny, and this trait was almost exclusively limited to Rhinocerotinae in our dataset (see Figure 1 and Figure S2). As outlined by Mitchell *et al.* [18], taxonomic scale can represent an important factor when interpreting CREA results.

In some cases, facial gracilisation can rather be observed at a finer taxonomic scale, due to shared ancestry and phylogenetic niche conservatism (i.e., tendency of closely related species to be more ecologically similar) [18]. Closely related species are more likely to share similar diets and biting requirements that may fit within a common allometric trajectory. Contrarily, eco-morphological shifts reflecting dietary specialisation coupled with highly modified cranial proportions that can contribute to the deviation from CREA may be more apparent at the broader phylogenetic scale [18]. Therefore, large-horned rhinos presumably showed facial gracilisation as a consequence of a similar plan of morphological organisation (i.e., morphotype) of their crania due to shared ancestry.

In summary, rhinoceroses join a few other mammalian groups that pose exceptions to CREA (e.g., hominins, sabre-toothed cats, horses), and therefore contribute to the understanding of this common evolutionary pattern [10,16,18,20,26]. The present work demonstrates that peculiar rostral adaptations, reflecting a series of ecomorphological shifts that affect the biomechanical and developmental properties of the cranium, can lead to divergence from predicted evolutionary trajectories. It should be acknowledged that the sample sizes for the subset test were relatively small, and that additional taxa might increase the power of statistical inference, potentially leading to the recognition of patterns not observed here. Taxonomic scale seemed to be an important factor in interpreting CREA results; therefore, a broader taxonomic sample should be included in future research to better understand evolutionary processes and drivers that have shaped rhinocerotid cranial diversity.

Results from this study, together with the recent work on craniofacial allometry in horses [26] provide important insight that CREA may be generally absent in perissodactyls, in contrast to ruminant artiodactyls, where it represents a ubiquitous pattern [23,27]. Rhoda *et al.* [27] hypothesised that the pervasiveness of CREA in ruminants may facilitate craniodental adaptations along the browser-grazer continuum, as it would allow simple shifts along the common allometric trajectory to accommodate cranial proportions in response to varying dietary pressures. Therefore, they suggested that the potential absence of CREA in perissodactyls (confirmed by this and the previous study [26]) may have provided artiodactyls an adaptive advantage throughout their radiation during the late Caenozoic. Although this hypothesis remains to be thoroughly explored in the future, present results and the mentioned studies offer a valuable starting point for its research.

### 4.3. Tempo of cranial evolution

As demonstrated by the present study, cranial adaptations associated with the horn presence did not noticeably influence the rate of morphological evolution, indicating that the cranial shape of hornless and horned rhinos had, on average, changed at the same tempo throughout their evolution. This suggests that the origin of such evolutionary novelties may not always be linked to a faster rate of morphological change, as previously shown in clades such as cetaceans [5] and sabre-toothed nimravids [6]. These modifications, resulting in the appearance of peculiar morphological traits, may rather occur at a more gradual, slower, or moderate tempo of evolution in some clades. Nevertheless, multiple pulses of increased evolutionary rate were recovered across rhinocerotid phylogeny.

The initial acceleration phase occurred in the early stage of Rhinocerotidae evolution during the Oligocene-Miocene transition (∼36.5-33.2 Mya). This burst is associated with the short internal branches of basal rhinocerotids that may reflect early rapid divergence, which would suggest a positive relationship between speciation rate and morphological evolution [123,124]. This morphological change may be related to the establishment of distinguishable Rhinocerotidae cranial morphology, including traits such as high zygomatic arches and a wide and deep nasal notch apparent with the emergence of *Epiaceratherium delemontense* [108]. Since the horn was not yet present in this early stage, this event emphasises the importance of traits unrelated to the horn variation in the evolution of Rhinocerotidae cranial morphology. Moderately high rates were inferred at the origin of two out of four major rhinocerotid clades, including elasmotheres and teleoceres, which were probably linked to the appearance of clade-specific morphological innovations. In elasmotheres, this rate shift may be related to the appearance of rostral cranial adaptations, since *Diceratherium* is considered the first rhinocerotid to develop the horn, leading to the evolution of more pronounced and robust nasal bones [62]. The second phase of rate acceleration, which occurred at the origin of teleoceres at the Oligocene-Miocene boundary (24-22.5 Mya), may be correlated with the evolution of multiple advanced traits in this group, such as brachycephalic skulls, more retracted nasal notches, or lower position of the infraorbital foramina [42]. However, it should be noted that teleoceres appeared intermixed with several acerathere species in our phylogeny, implying that the interpretation of this acceleration may change with further advances in understanding the relationship within Rhinocerotidae.

Throughout the Middle and Late Miocene (16-5.33 Mya), several phases of increased evolutionary rates were identified, especially within Rhinocerotinae (extant rhinos and their relatives). This geological period is marked by major climatic events of global cooling that took place after the warm Miocene climatic optimum (17 to 14 Mya), leading to the expansion of open grassland habitats [125–128]. This environmental transition has dramatically affected the evolution of many large herbivore clades and has prompted speciation and morphological diversification in multiple lineages (e.g., equids and bovids) [124,129–138]. In the present analysis, the most remarkable increase in the rate of cranial change was recorded during the evolution of the modern African rhinos. This acceleration phase started immediately upon the divergence of extant rhino species (∼14 Mya) and lasted until the latest Miocene (∼6 Mya). The peak took place from approximately 9-6 Mya, starting upon the divergence of African rhinos from Eurasian *Ceratotherium neumayri*. This whole acceleration phase closely overlaps with a transition from C3- to C4-dominated grassland habitats that began in the Early Miocene and further intensified throughout the Late Miocene, peaking around 7 to 5 Mya [127,128,132]. This is congruent with a slightly slower rate of cranial evolution in the *Ceratotherium*-*Diceros* ancestral lineage, followed by a rapid burst after the 9 Mya boundary that could be associated with an entering to a new adaptive zone within African grassland habitats.

Black (*D. bicornis*) and white rhinos (*C. simum*) occupy two opposite extremes of the plant diet spectrum, showing substantial cranial specialisation to browsing and grazing diets, respectively [43,139,140]. However, this divergent specialisation evolved after the split of the two species at the Miocene-Pliocene boundary (5 Mya), while their ancestors were considered to be mixed feeders that consumed a significant proportion of grass vegetation [63,64]. Therefore, we propose that an increased rate of morphological evolution in ancestral lineages of modern African rhinos likely reflects a shift in cranial shape linked to the adaptation to a more grazing diet, which assumed greater consumption of abrasive plant material. In line with this assumption, it has been shown that African rhinocerotids incorporated a substantial proportion of C4 grasses in their diets as early as 9.6 Mya, with this proportion rising significantly around 7 Mya and onwards [141], coinciding with the observed peak of cranial evolution. It is likely that cranial changes, linked to this dietary transition, represent modifications in the dental region to house larger, more hypsodont teeth or the shape of the nuchal crest to accommodate head position while grazing, enhancing the overall feeding efficiency associated with the utilisation of new resources.

In addition, a fast rate of cranial evolution was recovered upon divergence of the *Pliorhinus ringstroemi*-*Coelodonta antiquitatis* species pair from *Dicerorhinus sumatrensis*. This acceleration occurred from around 11.5 to 9.5 Mya and could be correlated with the acquisition of derived cranial traits present in *Pliorhinus* and *Coelodonta*, compared to the relatively unchanged morphology of *Dicerorhinus sumatrensis* [29,37,60], characterised by a low evolutionary rate. Moreover, a moderately high rate was observed in *C. antiquitatis*, which could also be associated with the evolution of cranial morphology adapted for a grazing diet in this species [142]. Certain herbivorous clades and species exhibited rapid changes in dental and other morphological traits during Neogene episodes of global cooling [15,124,133,136,143]. However, to our knowledge, the present results provide the first evidence of increased rates of cranial shape evolution during the transition from C3- to C4-dominated grasslands for any group of large herbivorous mammals, emphasising an important role of cranial adaptations in the rise of ecomorphological innovations under the pressure of global paleoclimatic events. The series of increased evolutionary rates within Rhinocerotinae may, in part, reflect their adaptability to changing environmental conditions during the Neogene and Quaternary, resulting in a spectrum of herbivore diets coupled with diverse cranial adaptations. These changes provide potential insight into the success of this clade, as they were the only rhino group to persist deeply into the Pliocene and Pleistocene, ultimately giving rise to all five extant species.

The observed pulses of accelerated morphological evolution across Rhinocerotidae appear to coincide with periods of global cooling, such as the Eocene-Oligocene transition, glaciation at the Oligocene-Miocene boundary, and the late Middle and Late Miocene [125,126,144–147]. These shifts in climate have triggered increased aridification and habitat heterogeneity [135,148,149], which potentially led to the utilisation of novel ecological opportunities in rhinocerotids, highlighting the importance of evolutionary novelties in shaping the tempo of morphological evolution across specific clades. Thus, extrinsic factors may have prompted cranial diversification in rhinocerotids through the series of niche shifts that reflect an increased rate of morphological innovation. A similar pattern of accelerated diversification during these global cooling events has already been documented for the evolution of inner ear shape in ruminants, another group of large herbivorous mammals with a broad spectrum of different ecologies [124]. However, to better understand the processes and mechanisms that have driven cranial evolution across Rhinocerotidae, direct assessment of the correlation between paleoclimatic events and morphological diversification is needed.

## 5. Conclusions

The present study highlights the role of horn diversification in the morphological evolution of Rhinocerotidae, demonstrating that cranial shape variation evolved in a phylogenetically constrained fashion. Rhinos contribute to the rising evidence of exceptions to the mammalian craniofacial evolutionary allometry (CREA) [10,18,26], most likely caused by a series of ecomorphological shifts across the clade that evolved independently of cranial size. These shifts primarily include rostral adaptations arising under selective pressure to accommodate different horn forms and dietary preferences, emphasising their general potential in inducing deviations from the expected allometric trajectory [10,18]. Multiple phases of accelerated morphological change were revealed as an important driver in generating the rhinos’ cranial diversity, providing insight into how major Caenozoic paleoclimatic events shaped the tempo of evolution in response to shifting environments, such as the transition from closed forest habitats to open grasslands. The present work sets the baseline for further research on the cranial evolution of rhinos, and future studies incorporating a broader taxonomic sample will be essential to fully elucidate the macroevolutionary dynamics of this remarkable yet endangered group of large herbivores.

## Acknowledgements

The authors would like to thank the Museo Regionale di Scienze Naturali di Torino (MRSN), Museo di Storia Naturale "La Specola" di Firenze (MZUF), Museo Civico di Zoologia di Roma (MCZR), and the Museo di Anatomia Comparata "B. Grassi" of Sapienza University for providing access to their collections and valuable specimens. We would also like to thank Muséum national d’Histoire naturelle (MNHN) for providing access to their digital collection of 3D models (3Dthèque), and we acknowledge the work of the following authors: Jérémy Tissier, Roger Benson, and Panagiotis Kampouridis. Animal silhouettes used for the visualisation were retained from PhyloPic (https://www.phylopic.org/), and we acknowledge the work of the following creators: Zimices, Kai Caspar, Steven Traver, Robert Bruce Horsfall, and Ivan Iofrida. Shillouetes were obtained under the CC0 1.0 Universal Public Domain Dedication licence, the CC BY-SA 3.0 Attribution-NonCommercial Unported licence, and the CC BY 4.0 Attribution International licence.

