## supplementary figures and tables for "Patterns and drivers of cranial evolution in rhinos (Rhinocerotoidea; Perissodactyla): allometry, phylogeny and tempo of evolution"

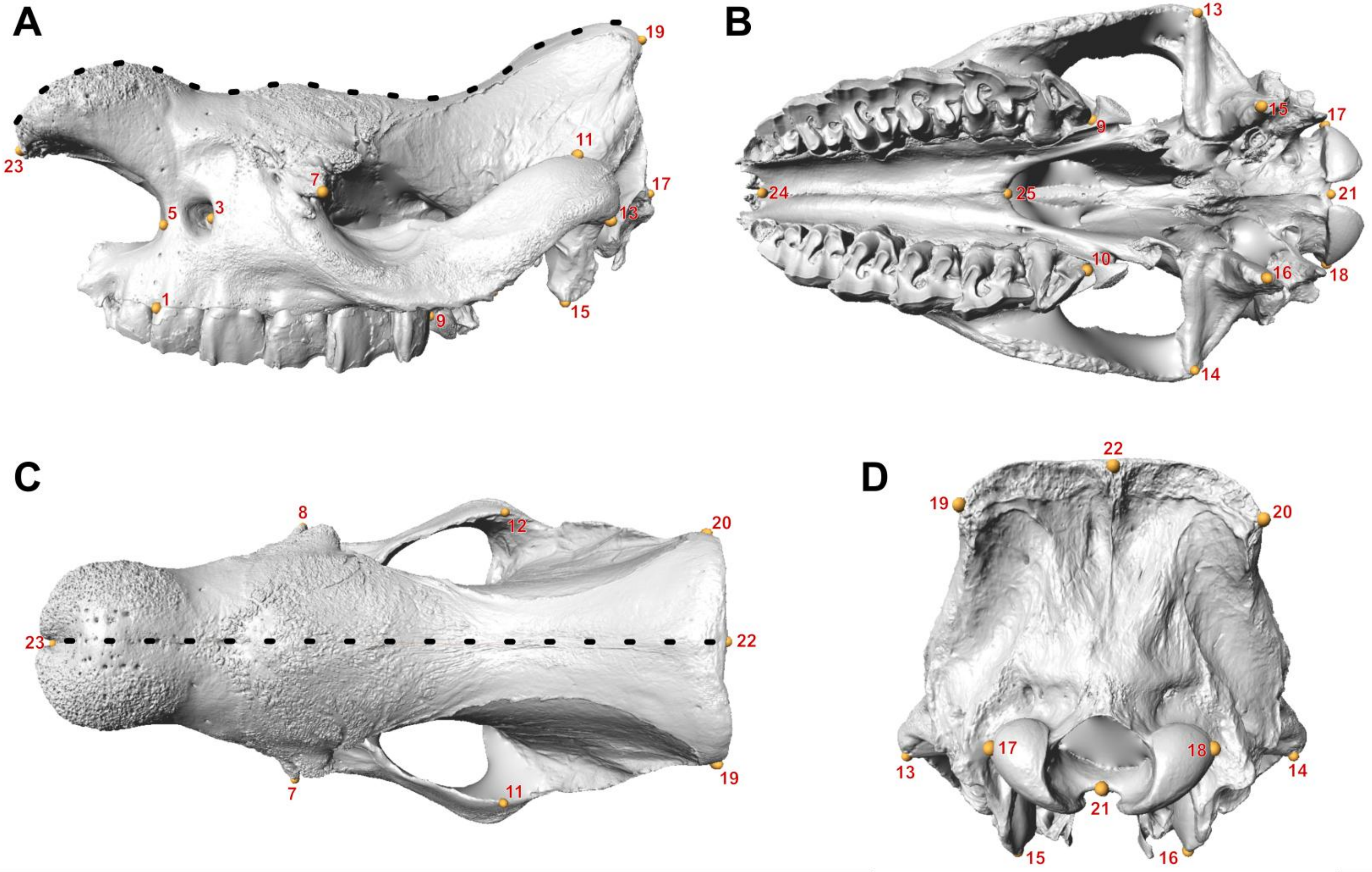


**Figure S1.** Landmark configuration used for geometric morphometric analysis of cranial morphology. Yellow points – fixed landmarks. Black points – semilandmarks. (A) Lateral view. (B) Ventral view. (C) Dorsal view. (D) Caudal view. The black rhino (*Diceros bicornis*) is shown as an example.

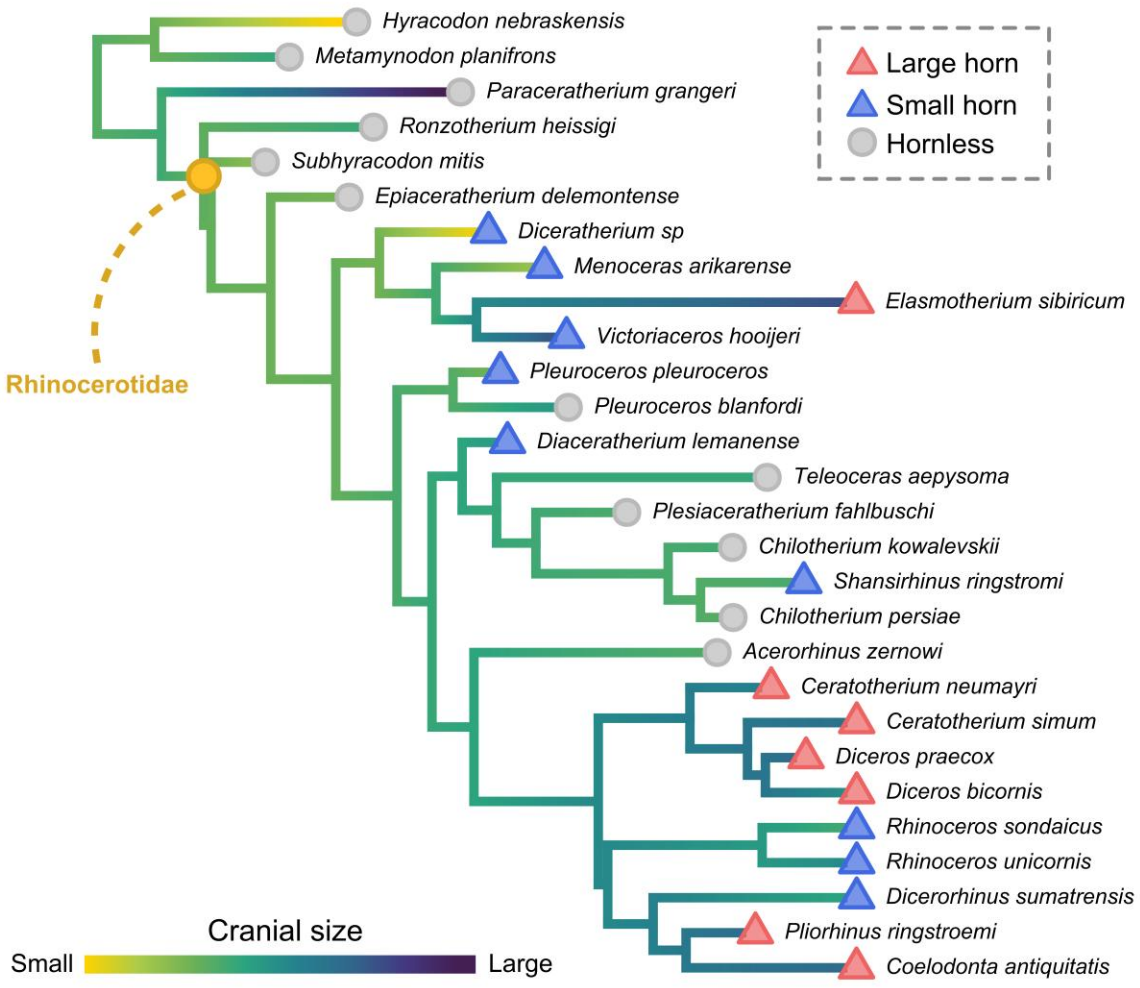


**Figure S2.** Ancestral state reconstruction of cranial size, with logarithm of centroid size mapped onto phylogeny. Symbols at the tree tips indicate presence of the horn within Rhinocerotoidea. Note that *Victoriaceros hooijeri* represents an exception as a large-sized rhinocerotid possessing a small horn.

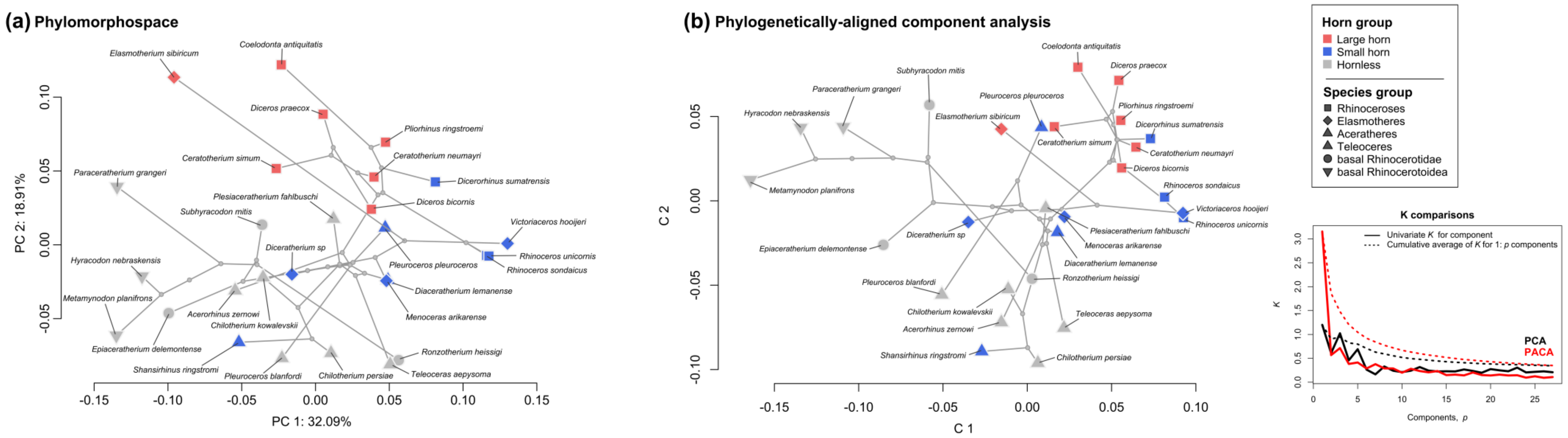


**Figure S3.** Phylomorphospace - standard PCA with mapped phylogeny (a). Phylogenetically-aligned component analysis (PACA) with comparison of phylogenetic signal (K) through *p* components (b). Note the similar distribution of taxa and horn groups between the two analyses (irrespective of some differences in data space rotation), indicating a strong influence of phylogeny on species distribution within the phylomorphospace.

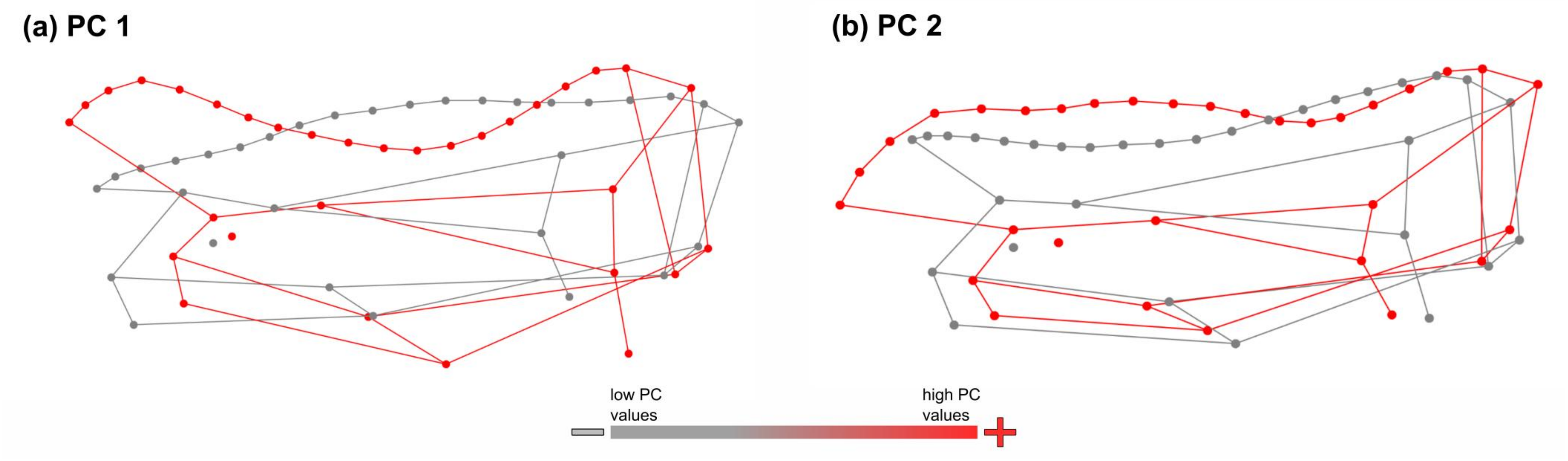


**Figure S4.** Cranial shape changes from the lateral view along the first two principal components from the phylomorphospace analysis, showing shapes at the two extremes.

**Table S1.** Table of contents including sources of fossil ages, and morphological and molecular data used in phylogenetic analysis.

| Species | Morphological data source | Fossil ages source | DNA accession IDs | Molecular data source |
|---|---|---|---|---|
| *Aceratherium incisivum* | Antoine et al. (2022) | NOW Database | / | / |
| *Acerorhinus zernowi* | Antoine et al. (2003) | NOW Database | / | / |
| *Alicornops alfambrense* | Antoine et al. (2003) | NOW Database | / | / |
| *Alicornops complanatum* | Antoine et al. (2003) | NOW Database | / | / |
| *Alicornops simorrense* | Antoine et al. (2022) | NOW Database | / | / |
| *Aprotodon fatehjangense* | Antoine et al. (2003) | NOW Database | / | / |
| *Brachypotherium brachypus* | Antoine et al. (2003) | NOW Database | / | / |
| *Brachypotherium perimense* | Antoine et al. (2022) | NOW Database | / | / |
| *Bugtirhinus praecursor* | Antoine et al. (2003) | NOW Database | / | / |
| *Ceratotherium neumayri* | Antoine et al. (2022) | NOW Database | / | / |
| *Ceratotherium simum* | Antoine et al. (2022) | extant | NC_001808.1 | Xu et al. (1996) |
| *Chilotherium anderssoni* | Antoine et al. (2003) | NOW Database | / | / |
| *Chilotherium kowalevskii* | Pandolfi (2015) | NOW Database | / | / |
| *Chilotherium perisae* | Pandolfi (2015) | NOW Database | / | / |
| *Coelodonta antiquitatis* | Antoine et al. (2022) | NOW Database | FJ905813.1 | Willerslev et al. (2009) |
| *Diaceratherium lemanense* | Antoine et al. (2003) | NOW Database | / | / |
| *Diceratherium* sp. | Antoine et al. (2003) | NOW Database | / | / |
| *Dicerorhinus fusuiensis* | Antoine et al. (2022) | NOW Database | / | / |
| *Dicerorhinus sumatrensis* | Antoine et al. (2022) | extant | FJ905816.1 | Willerslev et al. (2009) |
| *Diceros bicornis* | Antoine et al. (2022) | extant | FJ905814.1 | Willerslev et al. (2009) |
| *Diceros praecox* | Personal observations | NOW Database | / | / |
| *Dihoplus schleiermacheri* | Antoine et al. (2022) | NOW Database | / | / |
| *Dromoceratherium mirallesi* | Antoine et al. (2022) | NOW Database | / | / |
| *Elasmotherium sibiricum* | Antoine (2003) | Kosintsev et al. (2019) | MH937513.1 | Kosintsev et al. (2019) |

| | | | | |
|---|---|---|---|---|
| *Epiaceratherium bolcense* | Tissier et al. (2021) | NOW Database | / | / |
| *Epiaceratherium delemontense* | Tissier et al. (2021) | NOW Database | / | / |
| *Gaindatherium browni* | Antoine et al. (2022) | NOW Database | / | / |
| *Hispanotherium beonense* | Antoine et al. (2003) | NOW Database | / | / |
| *Hoploaceratherium tetradactylum* | Antoine et al. (2022) | NOW Database | / | / |
| *Hyrachyus eximius* | Antoine et al. (2022) | NOW Database | / | / |
| *Hyracodon nebraskensis* | Tissier et al. (2018) | NOW Database | / | / |
| *Lartetotherium sansaniense* | Antoine et al. (2022) | NOW Database | / | / |
| *Menoceras arikarense* | Antoine et al. (2003) | NOW Database | / | / |
| *Metamynodon planifrons* | Tissier et al. (2018) | NOW Database | / | / |
| *Nesorhinus hayasakai* | Antoine et al. (2022) | Antoine et al. (2022) | / | / |
| *Nesorhinus philippinensis* | Antoine et al. (2022) | Antoine et al. (2022) | / | / |
| *Paraceratherium bugtiense* | Tissier et al. (2018) | NOW Database | / | / |
| *Paraceratherium grangeri* | Tissier et al. (2018) | NOW Database | / | / |
| *Plesiaceratherium fahlbuschi* | Sun et al. (2024) | NOW Database | / | / |
| *Pleuroceros blanfordi* | Antoine et al. (2010) | NOW Database | / | / |
| *Pleuroceros pleuroceros* | Antoine et al. (2003) | NOW Database | / | / |
| *Pliorhinus megarhinus* | Antoine et al. (2022) | NOW Database | / | / |
| *Pliorhinus miguelcrusafonti* | Pandolfi et al. (2021) | NOW Database | / | / |
| *Pliorhinus ringstroemi* | Li et al. (2024) | NOW Database | / | / |
| *Prosantorhinus douvillei* | Antoine et al. (2003) | NOW Database | / | / |
| *Protaceratherium minutum* | Antoine et al. (2003) | NOW Database | / | / |
| *Rhinoceros kendengindicus* | Antoine et al. (2022) | Antoine et al. (2022) | / | / |
| *Rhinoceros platyrhinus* | Antoine et al. (2022) | NOW Database | / | / |
| *Rhinoceros sinensis* | Antoine et al. (2022) | NOW Database | / | / |
| *Rhinoceros sondaicus* | Antoine et al. (2022) | extant | FJ905815.1 | Willerslev et al. (2009) |

| | | | | |
|---|---|---|---|---|
| *Rhinoceros unicornis* | Antoine et al. (2022) | extant | NC_001779.1 | Xu et al. (1996) |
| *Ronzotherium filholi* | Tissiet et al. (2021) | NOW Database | / | / |
| *Ronzotherium heissigi* | Tissiet et al. (2021) | NOW Database | / | / |
| *Sellamynodon zimborensis* | Tissier et al. (2018) | Paleobiology Database | / | / |
| *Shansirhinus ringstromi* | Sun et al. (2024) | NOW Database; Deng (2005) | / | / |
| *Stephanorhinus etruscus* | Antoine et al. (2022) | NOW Database | / | / |
| *Stephanorhinus kirchbergensis* | Pandolfi et al. (2021) | NOW Database | KX646743.1 | Kirillova et al. (2017) |
| *Stephanorhinus pikermiensis* | Antoine et al. (2022) | NOW Database | / | / |
| *Subhyracodon mitis* | Antoine et al. (2003) | NOW Database | / | / |
| *Tapirus terrestris* | Antoine et al. (2022) | extant | AJ428947.1 | Janke et al. (unpublished) |
| *Teleoceras aepysoma* | Antoine et al. (2022) | NOW Database | / | / |
| *Trigonias osborni* | Antoine et al. (2022) | NOW Database | / | / |
| *Victoriaceros hooijeri* | Geraads et al. (2016) | NOW Database | / | / |

**Table S2.** Specimen table used for geometric morphometric analysis. Museum names and catalogue numbers are incorporated into IDs.

| ID | Species | Group | Scanned by/Source |
|---|---|---|---|
| Acerorhinus_zernowi_SNSB-BSPG_1968_6_203 | *Acerorhinus zernowi* | Aceratheres | Mihajlo Milić |
| Ceratotherium_neumayri_MNHN.PIK.971 | *Ceratotherium neumayri* | Rhinoceroses | 3Dthèque MNHN - Paris |
| Ceratotherium_simum_MRSN_7250 | *Ceratotherium simum* | Rhinoceroses | Naomi De Leo |
| Ceratotherium_simum_MZUF.C.7528 | *Ceratotherium simum* | Rhinoceroses | Naomi De Leo |
| Chilotherium_kowalevskii_SNSB-BSPG_1968_6_386 | *Chilotherium kowalevskii* | Aceratheres | Mihajlo Milić |
| Chilotherium_persiae_MNHN.MAR.3881 | *Chilotherium perisae* | Aceratheres | 3Dthèque MNHN - Paris |
| Coelodonta_antiquitatis_DMK.nocat | *Coelodonta antiquitatis* | Rhinoceroses | Sketchfab |
| Coelodonta_antiquitatis_MNHN.1926.4 | *Coelodonta antiquitatis* | Rhinoceroses | 3Dthèque MNHN - Paris |
| Diaceratherium_lemanense_MNHN.AC.2375.LIM.772 | *Diaceratherium lemanense* | Teleoceres | 3Dthèque MNHN - Paris |
| Diceratherium_sp_UW.4996 | *Diceratherium* sp. | Elasmotheres | Sketchfab |
| Dicerorhinus_sumatrensis_WML.1963.173.74 | *Dicerorhinus sumatrensis* | Rhinoceroses | Davide Tamagnini |
| Diceros_bicornis_AN.CO.111-360 | *Diceros bicornis* | Rhinoceroses | Naomi De Leo |
| Diceros_bicornis_MCZR_OS_0785 | *Diceros bicornis* | Rhinoceroses | Naomi De Leo |
| Diceros_bicornis_MRSN_nocat | *Diceros bicornis* | Rhinoceroses | Naomi De Leo |
| Diceros_praecox_KNM-ER.5555 | *Diceros praecox* | Rhinoceroses | African Fossils |
| Elasmotherium_sibiricum_NHMUK.M12429 | *Elasmotherium sibiricum* | Elasmotheres | Phenome10K |
| Epiaceratherium_delemontense_MJSN.POI.007–245 | *Epiaceratherium delemontense* | basal Rhinocerotidae | MorphoMuseuM |
| Hyracodon_nebraskensis_SNSB-BSPG_1988_1_74 | *Hyracodon nebraskensis* | basal Rhinocerotoidea | Mihajlo Milić |
| Menoceras_arikarense_UF.VP.455000 | *Menoceras arikarense* | Elasmotheres | Sketchfab |
| Metamynodon_planifrons_UNISTRA.2015.0.1106 | *Metamynodon planifrons* | basal Rhinocerotoidea | MorphoMuseuM |

| | | | |
|---|---|---|---|
| Paraceratherium_grangeri_NMHUK.M84196 | *Paraceratherium grangeri* | basal Rhinocerotoidea | Phenome10K |
| Plesiaceratherium_fahlbuschi_SNSB-BSPG_1959_2_400 | *Plesiaceratherium fahlbuschi* | Aceratheres | Mihajlo Milić |
| Pleuroceros_blanfordi_SNSB-BSPG_1956_2 | *Pleuroceros blanfordi* | Aceratheres | Mihajlo Milić |
| Pleuroceros_pleuroceros_MNHN.LIM.778 | *Pleuroceros pleuroceros* | Aceratheres | 3Dthèque MNHN - Paris |
| Pliorhinus_ringstroemi_SNSB-BSPG_2000_1_56 | *Pliorhinus ringstroemi* | Rhinoceroses | Mihajlo Milić |
| Rhinoceros_sondaicus_MRSN_694 | *Rhinoceros sondaicus* | Rhinoceroses | Naomi De Leo |
| Rhinoceros_sondaicus_WML.1988.214.71 | *Rhinoceros sondaicus* | Rhinoceroses | Davide Tamagnini |
| Rhinoceros_unicornis_A.8085-Dotta.ACM.4252 | *Rhinoceros unicornis* | Rhinoceroses | Davide Tamagnini |
| Ronzotherium_heissigi_MNHN.LIM.181 | *Ronzotherium heissigi* | basal Rhinocerotidae | 3Dthèque MNHN - Paris |
| Shansirhinus_ringstromi_SNSB-BSPG_2002_1_1 | *Shansirhinus ringstromi* | Aceratheres | Mihajlo Milić |
| Subhyracodon_mitis_YPM.VP.010253 | *Subhyracodon mitis* | basal Rhinocerotidae | Phenome10K |
| Teleoceras_aepysoma_ETMNH.609 | *Teleoceras aepysoma* | Teleoceres | Sketchfab |
| Victoriaceros_hooijeri_KNM-KA.57652 | *Victoriaceros hooijeri* | Elasmotheres | 10.1080/02724634.2016.1103247 |

**Table S3.** Definitions of fixed landmarks used for geometric morphometrics.

| Landmark | Description |
|---|---|
| 1, 2 | Antero-lateral point on the alveolar margin of the third premolar |
| 3, 4 | Center of the infraorbital foramen |
| 5, 6 | Deepest (most posterior) point of the nasal notch |
| 7, 8 | Tip of the caudal lacrimal process |
| 9, 10 | Most posterior point of the tooth row |
| 11, 12 | The highest point of the zygomatic arch |
| 13, 14 | Most lateral point of the articular tubercle |
| 15, 16 | Ventral tip of the postglenoid process |
| 17, 18 | Most lateral point of the occipital condyle |
| 19, 20 | Most lateral point of the dorsal part of the nuchal crest |
| 21 | Basion: most anterior point of the foramen magnum |
| 22 | Inion: most posterior point of the cranium |
| 23 | Rhinion: most anterior midline point on the nasals |
| 24 | Most posterior point of the foramen incisivum |
| 25 | Most posterior point of the palatine along the midline |
